# The magnitude of the secondary B cell response is primarily defined by antibody feedback inhibition rather than the number of memory B cells present

**DOI:** 10.64898/2026.02.20.707093

**Authors:** Malte J. Hoormann, Noémi Becza, Lingling Yao, Anton V. Gorbachev, Stefanie Kuerten, Magdalena Tary-Lehmann, Greg A. Kirchenbaum, Paul V. Lehmann

**Author notes:** These authors contributed equally to this work and share senior authorship (Greg A. Kirchenbaum, Paul V. Lehmann). Author to whom correspondence should be addressed (G.A.K). (M.J.H.); (N.B.); (L.Y.); (P.V.L.), (S.K.), (A.G); (MTL).

## Abstract

Clonal expansion of memory lymphocytes after each antigen encounter is the primary mechanism for amplifying immunity. For most vaccines, boosters are common practice and are expected to stimulate proliferation of pre-existing memory B cells (B_mem_), thereby expanding the antigen-specific B_mem_ pool, along with driving their differentiation into antibody-producing plasma cells that replenish antibody titers. It is widely assumed that the number of B_mem_ present in the body prior to administration of a booster vaccination will define the magnitude of the ensuing response. However, due to technical limitations hampering reliable detection of rare antigen-specific B_mem_ in human subjects, the extent to which B_mem_ numbers are actually modulated following a booster vaccination remains unclear. By comparing B_mem_ frequencies and antibody titers in the same individuals after primary and secondary vaccination with SARS-CoV-2 Spike (S-antigen)-encoding mRNA we found that expansion of B_mem_ and the magnitude of the secondary antibody response were not determined by the number of B_mem_ measured before the second vaccine inoculation. Instead, both were inversely correlated with levels of S-antigen-specific serum IgG prior to the secondary antigen exposure. Collectively, the data suggest that secondary B cell responses are constrained by antibody feedback inhibition of B_mem_, rather than their paucity.

## 1. Introduction

Humoral immunity is mediated by the B cell system. When a primary B cell response is engaged after natural infection or the first administration of a vaccine, naïve antigen-specific B cells are recruited and become activated. The frequency of such naïve antigen-specific B cells is very low in individuals who have not previously encountered the antigen – it is estimated to be as low as 1 in 10^11^ of all B cells in the body (1, 2). During the primary immune response, these naïve B cells proliferate and differentiate into antibody-secreting plasma cells (PC) or memory B cells (B_mem_). Owing to their increased frequency, B_mem_ enable rapid secondary recall responses through their ability to undergo additional waves of proliferation – their progeny giving rise to both PC and B_mem_ – upon re-encountering the same or related antigen (3).

Vaccination-induced immune responses in humans and experimental animals have been extensively studied, primarily relying on antibody measurements. Recent data suggest, however, that antibody titers are a poor reflection of the underlying B cell reactions. Murine models revealed that divergent differentiation pathways define the generation of PC and B_mem_ during the antigen-driven B cell response (4–8). As the precursor B cells (naïve cells in primary or B_mem_ in secondary responses) proliferate in germinal centers, the antigen (B cell) receptors (BCRs) of their daughter cells undergo somatic hypermutation, and as a consequence, subclones arise with a spectrum of different affinities for the triggering antigen (9). Of these subclones, only those with the highest affinity for the eliciting antigen (the homotype) are selected for PC differentiation, i.e., antibody production, whereas subclones with lower BCR affinity for the homotypic antigen exit lymphoid tissues as B_mem_. The biological importance of this divergent differentiation of PC and B_mem_ lineages is not only that it serves to increase antibody affinity for the homotype (affinity maturation), but it also facilitates the generation and retention of a diversified B_mem_ BCR repertoire that is poised to respond to future mutants of the original virus strain (heterotypes) if escape from neutralizing antibodies induced by the homotype occurs (10). It is also important to note that while both PC and B_mem_ can be long-lived, the lifespan of PC can be limited by competition for “niches” in the bone marrow, whereas B_mem_ do not experience such limitations (11–15). Furthermore, it is also important to consider that – whereas antibodies are extremely robust in vitro – in vivo, their half-life is relatively short, being about 3 weeks for IgG (16, 17).

Based on the above understanding of B cell immunobiology, it can be anticipated that serum antibodies and B_mem_ provide independent and different information regarding an individual’s immune status. Antibodies provide a snapshot of a waning immune response resulting from prior antigen encounters and constitute the “first wall of humoral immune defense” against the next encounter, whereas B_mem_ comprise the cellular basis for initiating a rapid and flexible secondary B cell response upon re-exposure to the homotypic- or heterotypic antigen, acting as the “second wall of humoral immunity” (3, 10, 18).

Measuring the antibody response following vaccination is the gold standard for human immune monitoring because techniques for quantifying antigen-specific immunoglobulins (Ig) in solution are well established, and because Ig are robust molecules after their isolation from the body (19). In contrast, the cellular basis of the antibody response is largely understudied due to the paucity of suitable test systems and the fragility of the cells to be interrogated. Most of our understanding of the cellular basis of immune responses comes from elegant but simplistic and short-term murine models involving atypically small model antigens. The detection of antigen-specific B_mem_ in mice and humans has been challenging primarily due to the very low frequency at which B_mem_ occur in the lymphocyte pool, even after clonal expansion (20) whereby in humans, frequencies of <<1 in 10^5^ among peripheral blood mononuclear cells (PBMC) are common even for B_mem_ specific for endemic viruses such as EBV, HCMV, and seasonal influenza viruses (21).

Over the last few years, two techniques have emerged as candidates for monitoring rare antigen-specific B_mem_ in peripheral blood. One method is staining B cells in PBMC with fluorescence-labeled antigen probes followed by their detection using flow cytometry (FC). The second is B cell ELISPOT/FluoroSpot (here collectively designated “ImmunoSpot”). In ImmunoSpot assays, the antigen-specific B cells are visualized via their antibody-derived secretory footprints and are quantified as spot-forming units (SFUs) (See Suppl. Figure S1). A recent direct side-by-side comparison (22) showed that FC and ImmunoSpot both detected SARS-CoV-2 Spike (S-) antigen-specific B_mem_ with comparable sensitivity and diagnostic specificity in PBMC provided that the B_mem_ were present in the mid- to high frequency range. Due to the inherent background noise of FC (20, 23) that is virtually absent in B cell ImmunoSpot assays, we hypothesized – and have proven – that the ImmunoSpot methodology is suited for the detection of very rare antigen-specific B_mem_ even in the <1 in 10^6^ PBMC frequency range (24). Additionally, the substantially lower volume of blood (or number of PBMC) required for ImmunoSpot analysis (21) prompted our choice of this platform for dissecting the cellular basis of the B_mem_ response in this study.

The B cell ELISPOT assay methodology was first reported over forty years ago (25, 26) and was then later adapted for performance in 96 well plates (27, 28). Notably, in first-generation B cell ELISPOT assays the nature of antigens that were suited for direct coating onto the test plate was more limited and did not permit reliable detection of antigen-specific secretory footprints for many (mostly hydrophilic) antigens; only after we introduced the affinity capture coating strategy (29, 30) (illustrated in Suppl. Fig. S1A) did this test system become universally applicable for efficient and high density coating of any antigen. In particular, this permitted us to reveal secretory footprints originating from individual SARS-CoV-2 S-antigen-specific B_mem_. Here, we studied secondary B_mem_ expansions and their dependence on B_mem_ frequencies and specific serum antibody titers present in the same individuals prior to booster vaccination.

Due to the paucity of data on the cellular basis of B cell responses, and in spite of incidental reports (31), it remains an open question whether pre-existing specific serum antibodies positively or negatively influence B_mem_ expansion and maturation following antigen re-encounter, i.e., whether they enhance or suppress the secondary B cell response. Pre-existing specific antibodies have been implicated in both immune stimulatory and immune suppressive effects (reviewed in (32, 33) and summarized in Suppl. Figure S2). Stimulatory antibody feedback regulation has been attributed to antigen deposition on follicular dendritic cells (FDCs) (34–38), complement receptor 2 stimulation (32, 39, 40), and antigen uptake by dendritic cells (DCs) (33, 41–44); suppressive effects have been tied to epitope masking (31, 45–49), B cell inhibition via the FcγR (32, 50–52), and antigen clearance (32, 33, 53). Presently, it is unclear which, if any, of these mechanisms prevail during a prime-boost immunization, and in particular how the secondary expansion of B_mem_ is affected by pre-existing antibody titers (31, 54, 55). Our studies of B_mem_ in parallel with serum antibodies aimed at gaining insights into this fundamental question; which has translational relevance to the development of optimal booster vaccination protocols.

For several reasons, we selected the SARS-CoV-2 “model” to study B_mem_ development after boosting. SARS-CoV-2 is unique among human viruses in that there is a defined timepoint at which the virus appeared and started to spread in the United States early in 2020 (56–58). Second, there appears to be no significant B cell cross-reactivity between the (Spike) S-antigen expressed by previously circulating common cold coronaviruses and SARS-CoV-2 (Yao et al, manuscript in preparation), suggesting that a truly naïve B cell population was engaged following the first vaccination. Moreover, the initial infections with SARS-CoV-2 were monitored in the population to a degree never before achieved for other pathogens. The individuals we tested were vaccinated soon after COVID-19 mRNA vaccines, encoding the Wuhan-Hu-1 strain of S-antigen, became available in 2020. Additionally, we also verified that each test subject had not been infected with SARS-CoV-2 before or during the observation period. In this way, we ensured that the induction of genuinely primary and secondary B cell responses in previously naïve people could be studied without pre-existing B cell memory or SARS-CoV-2 infection(s) complicating the interpretation of the results.

## 2. Materials and Methods

### 2.1 Human Samples

Blood samples were collected internally at Cellular Technology Limited (CTL) under an Advarra approved IRB #Pro00043178 (CTL contract laboratory study number GL20-16 entitled COVID-19 Immune Response Evaluation). Blood draws were collected at three timepoints: pre-bleed (P), 14 days after the first COVID-19 mRNA vaccination (D1), and 14 days following the second COVID-19 mRNA booster vaccination (D2) (Figure 1A). Blood was collected into either sodium heparin or serum/SST tubes. Peripheral blood mononuclear cells (PBMC) were isolated by standard density centrifugation and cryopreserved using serum free cryopreservation reagents (CTL Reagents LLC, Shaker Heights, Ohio, USA) according to previously described protocols (59). PBMC samples were stored in vapor nitrogen tanks until testing. For isolation of serum, blood was kept at room temperature for at least 1h before separation by centrifugation. Serum was transferred to cryovials and subsequently stored at −20°C until testing. Details of all human donors in this manuscript, including demographics, collection dates and vaccine manufacturer, are provided in Suppl. Table S1.

### 2.2 Polyclonal B cell Stimulation

Thawing, washing and counting of PBMC were performed as previously described (60, 61). Briefly, cells were transferred into polyclonal B cell stimulation cultures within 2 h of thawing after resuspension in complete medium (CM) comprising RPMI 1640 (Alkali Scientific, Fort Lauderdale, FL) supplemented with 10% fetal bovine serum (Gemini Bioproducts, West Sacramento, CA), 100 U/mL penicillin, 100 U/mL streptomycin, 2 mM L-Glutamine, 1 mM sodium pyruvate, 8 mM HEPES (all from Life Technologies, Grand Island, NY) and 50 µM β-mercaptoethanol (Sigma-Aldrich, St. Louis, MO). Polyclonal stimulation of donor PBMC was performed with Human B-Poly-S (CTL), containing the TLR7/8 agonist R848 and recombinant human IL-2 (62), at 0.5-2 x 10^6^ cells/mL in 25 cm^2^ tissue culture flasks (Corning, Sigma-Aldrich). PBMC were cultured at 37°C, 5% CO_2_ for five days to promote terminal differentiation of resting B cells into antibody-secreting cells (ASCs) prior to evaluation in ImmunoSpot® assays as described previously (63, 64).

### 2.3. Recombinant Proteins

Recombinant full-length SARS-CoV-2 Spike (S-antigen) protein representing the ancestral Wuhan-Hu-1 strain (65) was acquired from the Center for Vaccines and Immunology (CVI) (University of Georgia (UGA), Athens, GA, USA). Recombinant SARS-CoV-2 Nucleocapsid (NCAP) protein was purchased from the Wuhu Interferon Biological Products Industry Research Institute (Wuhu, China). Importantly, both recombinant proteins used in this study possessed a genetically-encoded His affinity tag enabling their affinity coating to ImmunoSpot® membrane plates.

**Figure 1.**
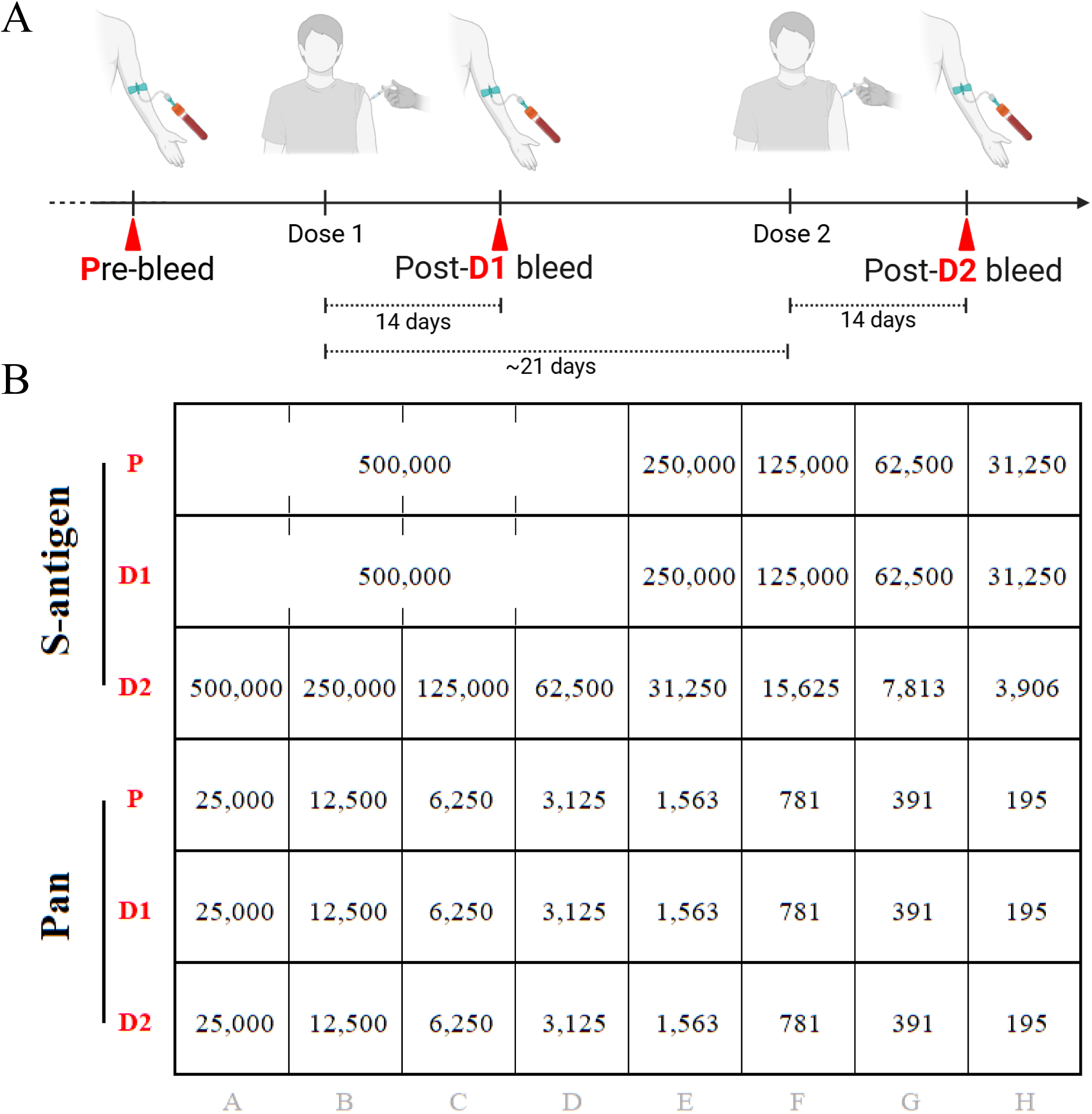
Study rationale. (A) Schematics of the vaccination regimen and blood draws (created with BioRender). The time interval between dose 1 (D1) and dose 2 (D2) vaccinations was according to the FDA-recommended schedule, ∼3 weeks apart. (B) ImmunoSpot® plate layout permitting S-antigen-specific IgG^+^ memory B cell (B_mem_) frequency measurements over an extended range. Pre-vaccination (designated Pre-bleed, P) and post-D1 samples were seeded at 5 x 10^5^ PBMC into four replicate wells; which permitted enumeration of individual S-antigen-reactive B_mem_ within a total of 2 x 10^6^ PBMC for the detection of low frequency events. Additionally, a two-fold serial cell dilution was also performed to extend the ability for precise counting when the S-antigen-specific secretory footprints were too crowded at the 5 x 10^5^ cells per well input for reliable enumeration. Moreover, because S-antigen-specific IgG^+^ B_mem_ frequencies were expected to be higher after the booster vaccination, post-D2 samples were tested using a two-fold serial dilution approach starting at 5 x 10^5^ live cells per well, yielding 8 titration points.

### 2.3 Multiplexed antigen-specific ImmunoSpot® assays with affinity capture coating

Antigen-specific ASCs were detected in multiplexed ImmunoSpot® assays using the affinity capture coating method (29). Low autofluorescence ImmunoSpot® assay plates were pre-conditioned with 70% (*v/v*) EtOH (15 μL/well), followed by two washes with phosphate-buffered saline (PBS) (180 μL/well). Assay plates were then coated with purified anti-His antibody at 10 µg/mL in Diluent A (provided in CTL’s affinity coating kits) overnight at 4°C. The follow day, ImmunoSpot® assays plates were washed once with 180 μL/well PBS and then coated overnight at 4°C with His-tag-labeled recombinant S-antigen or NCAP protein at 10 µg/mL in Diluent A. Prior to the addition of PBMC, ImmunoSpot® assay plates were washed once with PBS and then wells were blocked with CM for 1 h at room temperature (RT). Immediately prior to plating of PBMC, ImmunoSpot® assay plates were decanted and 100 μL pre-warmed CM was added to each well.

PBMC were harvested after 5 days of polyclonal stimulation and washed twice with PBS. PBMC were counted using CTL’s Live/Dead Cell Counting Suite on an ImmunoSpot^®^ S6 Flex Analyzer (CTL Analyzers LLC). Cell pellets were resuspended at 5 × 10^6^ live cells/mL in CM and used immediately in ImmunoSpot^®^ assays unless otherwise indicated.

To enhance the sensitivity of the S-antigen-specific ImmunoSpot® assay for measurements of rare antigen-reactive ASCs, pre-bleed (P) and post-D1 samples were tested in four replicate wells seeded with 5 x 10^5^ polyclonally stimulated PBMC, followed by a two-fold serially dilution in single wells providing four additional cell titration points. Since frequencies of S-antigen-specific ASCs were expected to be higher following the second (booster vaccination) antigen encounter, post-D2 PBMC samples were tested using a singlet two-fold serial dilution approach starting at 5 x 10^5^ PBMC per well, yielding 8 titration points. To avoid damage to the ImmunoSpot® assay plate, serial dilutions were performed in round bottom 96-well tissue culture plates (Corning, Sigma-Aldrich) and PBMC were subsequently transferred into the ImmunoSpot® assay plates, as previously described (64). Alternatively, for detection of NCAP-reactive ASC, PBMC samples were tested in four replicate wells seeded with 5 x 10^5^ polyclonally stimulated PBMC; however, owing to limited cell material some samples were only tested in two or three replicate wells. Following the addition of PBMC, ImmunoSpot® assay plates were incubated for 16 h at 37°C, 5% CO_2_. Plate-bound spot-forming units (SFUs), representing the secretory footprint of individual ASCs, were visualized using detection reagents included in the human IgG1/IgG2/IgG3/IgG4 Four-Color ImmunoSpot® kit (from CTL) according to the manufacturer’s instructions.

### 2.4. Multiplexed pan IgG subclass detection

Pan (total) IgG subclass (IgG1/IgG2/IgG3/IgG4) Four-Color ImmunoSpot® assays were performed in parallel with the antigen-specific assays described above to confirm ASC functionality in the polyclonally stimulated PBMC samples. To detect all IgG^+^ ASCs irrespective of their antigen specificity, cell suspensions were serially diluted two-fold in singlet starting at 2.5 × 10^4^ PBMC per well. Cell dilutions were performed in round bottom 96-well tissue culture plates as described above and PBMC were subsequently transferred into low autofluorescence ImmunoSpot® assay plates that were previously coated overnight at 4°C (following pre-wetting with EtOH as described above) with anti-κ/λ capture antibody provided in the human IgG1/IgG2/IgG3/IgG4 Four-Color ImmunoSpot^®^ kit (from CTL). The following day these ImmunoSpot® assay plates were washed and blocked as described above. After transfer of PBMC, assay plates were incubated for 16 h at 37°C, 5% CO_2_. Plate-bound SFUs representing the secretory footprints of individual IgG+ ASC were subsequently visualized using detection reagents included in the human IgG1/IgG2/IgG3/IgG4 Four-Color ImmunoSpot® kit (from CTL) according to the manufacturer’s instructions.

### 2.4. ImmunoSpot® Image Acquisition and SFU Counting

ImmunoSpot® plates were air-dried prior to scanning and counting on an ImmunoSpot® Ultimate S6 Analyzer using the Fluoro-X suite of ImmunoSpot® software (Version 7.0.28) and the Basic Count mode (CTL). Individual well images were quality controlled as needed to remove artifacts and to improve the accuracy of counts. Only SFU counts within the linear titration range of the ImmunoSpot® assay, or those from the highest cell input tested, were considered for frequency calculations and were subsequently used to extrapolate SFU counts to a fixed input of 10^6^ PBMC. As ImmunoSpot® multi-color B cell kits, analyzers, and software proprietary to CTL were used in this study, we refer to the collective methodology as ImmunoSpot^®^. Notably, since only IgG1^+^ and IgG3^+^ SFU were detected in S-antigen-specific ImmunoSpot® assays the resulting counts were combined and are denoted as IgG.

### 2.5. ELISA

S-antigen-specific IgG levels in serum samples were quantified according to previously described methods with minor modifications (66). In brief, microtiter ELISA plates (Immulon® 4HBX, flat-bottom, Thermo Fisher Scientific, Waltham, MA, USA) were coated with 80 μl/well of S-antigen at 2 μg/mL in carbonate buffer at pH 9.3 overnight at 4°C. Plates were then decanted and blocked with ELISA blocking buffer consisting of PBS supplemented with 2% (w/v) bovine serum albumin (Sigma-Aldrich, St. Louis, MO, USA) and 0.1% (v/v) Tween20 for 1.5 h at RT. After blocking, serially diluted serum samples were incubated in the microtiter plates overnight at 4°C. The following day, microtiter plates were washed with PBS prior to the addition of horseradish peroxidase (HRP)-conjugated anti-human IgG detection reagent (from CTL) and incubation for 1.5 h at RT. After washing the microtiter plates with PBS, 100 µL/well of 1-Step™ TMB ELISA Substrate Solution (Thermo Fisher Scientific) was added to develop the assay. Conversion of the TMB substrate was terminated by addition of 100 µL/well of 2 M HCl, and the optical density was measured at 450 nm using a Spectra Max 190 plate reader (Molecular Devices, San Jose, CA, USA). The abundance of S-antigen-specific IgG in the serum samples was determined by interpolating OD_450_ values into μg/mL of IgG equivalents using SpotStat^TM^ (Version 1.6.6.0, CTL). The underlying standard curves were generated by directly coating decreasing quantities of a reference IgG preparation (from Athens Research and Technology, Athens, GA, USA) in duplicates into designated wells of each assay plate.

### 2.4. Statistical Methods

Statistically significant differences in the frequencies of S-antigen-specific IgG^+^ ASCs and serum IgG levels between post-D1 and post-D2 timepoints were assessed using paired t-tests (GraphPad Prism 10, Version 10.6.1; San Diego, CA, USA) and are indicated in the corresponding figure legends. Pearson correlation and simple linear regression analyses were performed to evaluate associations between antigen-specific ASC frequencies and serum IgG levels at matching study timepoints. In the same manner, correlations were assessed between S-antigen-specific ASC frequencies at the post-D1 and post-D2 bleeds, between post-D1 bleed S-antigen-specific ASC frequencies and post-D2 bleed serum IgG levels, and between post-D1 and post-D2 bleed serum IgG levels, respectively. R^2^ values, 95% confidence intervals, and p-values are shown in the respective figures.

## 3. Results and Discussion

### 3.1. Overall Rationale

The timeline of sample collection in our study and the plate layout of the SARS-CoV-2 Spike (S-antigen)-specific ImmunoSpot assays that were performed are depicted in Figure 1. In essence, we focused on determining S-antigen-specific memory B cell (B_mem_) frequencies and antibody titers before (pre-bleed, P), 14 days after the first dose (post-D1), and 14 days after the second dose (post-D2) of COVID-19 mRNA vaccine (Comirnaty®) encoding S-antigen of the prototype SARS-CoV-2 (Wuhan-Hu-1 strain) virus. B cell reactivity to the SARS-CoV-2 Nucleocapsid (NCAP) protein, which was not included in the vaccine, was also measured to control for the possibility that infection with SARS-CoV-2 occurred prior to, or during, the observation period. One individual did indeed become positive for NCAP-specific B_mem_-derived IgG^+^ antibody-secreting cell (ASC) reactivity at the post-D2 timepoint and was therefore excluded from the analysis.

The overall goal of this study was to determine whether secondary clonal expansions of B_mem_ were preprogrammed when the antigen is re-encountered or are under feedback regulation. If preprogrammed, one would expect that individuals exhibiting higher frequencies of B_mem_ after the primary vaccination and therefore possessing increased numbers of precursor cells available for the secondary antigen encounter, would develop higher numbers of B_mem_ following renewed clonal expansion. In contrast, in the case of antibody feedback inhibition, frequencies of S-antigen-specific B_mem_ following the booster vaccination would be inversely related to the titer of Spike-specific IgG in the vaccinees at the post-D1 timepoint.

### 3.2. Divergence of B_mem_ frequencies and serum antibody titers

Antigen-specific B_mem_ can occur over a very wide frequency range in peripheral blood mononuclear cells (PBMC) (21, 67). The plate layout we used for ImmunoSpot® testing accommodated such differences (Figure 1B). The PBMC of each donor were plated at 5 x 10^5^ cells/well into four replicate wells, followed by additional two-fold serial dilutions of the sample, tested as singlets. This layout sets the lower limit of detection at one S-antigen-specific B_mem_-derived IgG^+^ ASC in 2 x 10^6^ PBMC, while the serial dilution permits the precise enumeration of B_mem_ occurring at higher frequencies (24). From the linear range of the counts of spot-forming unit (SFU) the frequency of antigen-specific B_mem_-derived IgG^+^ ASC can be extrapolated from single well serial dilutions with nearly identical precision as tests employing replicate wells for each cell dilution (21). In this manner, ImmunoSpot® assays were performed to establish the frequency of S-antigen-specific B_mem_-derived ASC secreting each of the four IgG subclasses. Additionally, in parallel, we also evaluated the PBMC samples in pan (total) IgG ImmunoSpot® assays to determine the frequency of ASC capable of secreting the different IgG subclasses irrespective of their antigen specificity.

Original well images depicting the testing of a representative subject (Donor 16) for S-antigen-specific B_mem_-derived IgG1^+^ ASC at the three PBMC collection timepoints is shown in Figure 2A and 2B, while the results for all donors are summarized in Figure 3A. Notably, owing to the elevated frequency of S-antigen-specific IgG^+^ B_mem_ detected at in post-D2 timepoint, serial dilution of the Donor 16-D2 sample permitted an accurate frequency measurement (Figure 2C). Moreover, no S-antigen-reactive IgG-producing B_mem_ were detected at the pre-bleed timepoint among the 2 x 10^6^ PBMC tested per sample, consistent with the notion that such donors were immunologically naïve to the SARS-CoV-2 S-antigen prior to their first vaccination (24). S-antigen-specific IgG^+^ B_mem_ became detectable in all subjects by day 14. As expected, considerable inter-individual B_mem_ frequency differences were seen in this cohort, ranging from 4- to 184 IgG^+^ ASC/10^6^ among the PBMC (Figure 3A). These data suggest that the magnitudes of initial clonal expansions differed significantly between these individuals. Moreover, in light of recent observations (48), it is tempting to postulate that the primary B cell response is also shaped by antibody feedback inhibition and that this feedback would be most accentuated in individuals who more rapidly mounted an antibody response. To this end, quantification of specific IgG serum antibody levels also showed a relatively high degree of heterogeneity (Figure 3B), as typically observed for post-vaccination antibody responses (68–74).

**Figure 2.**
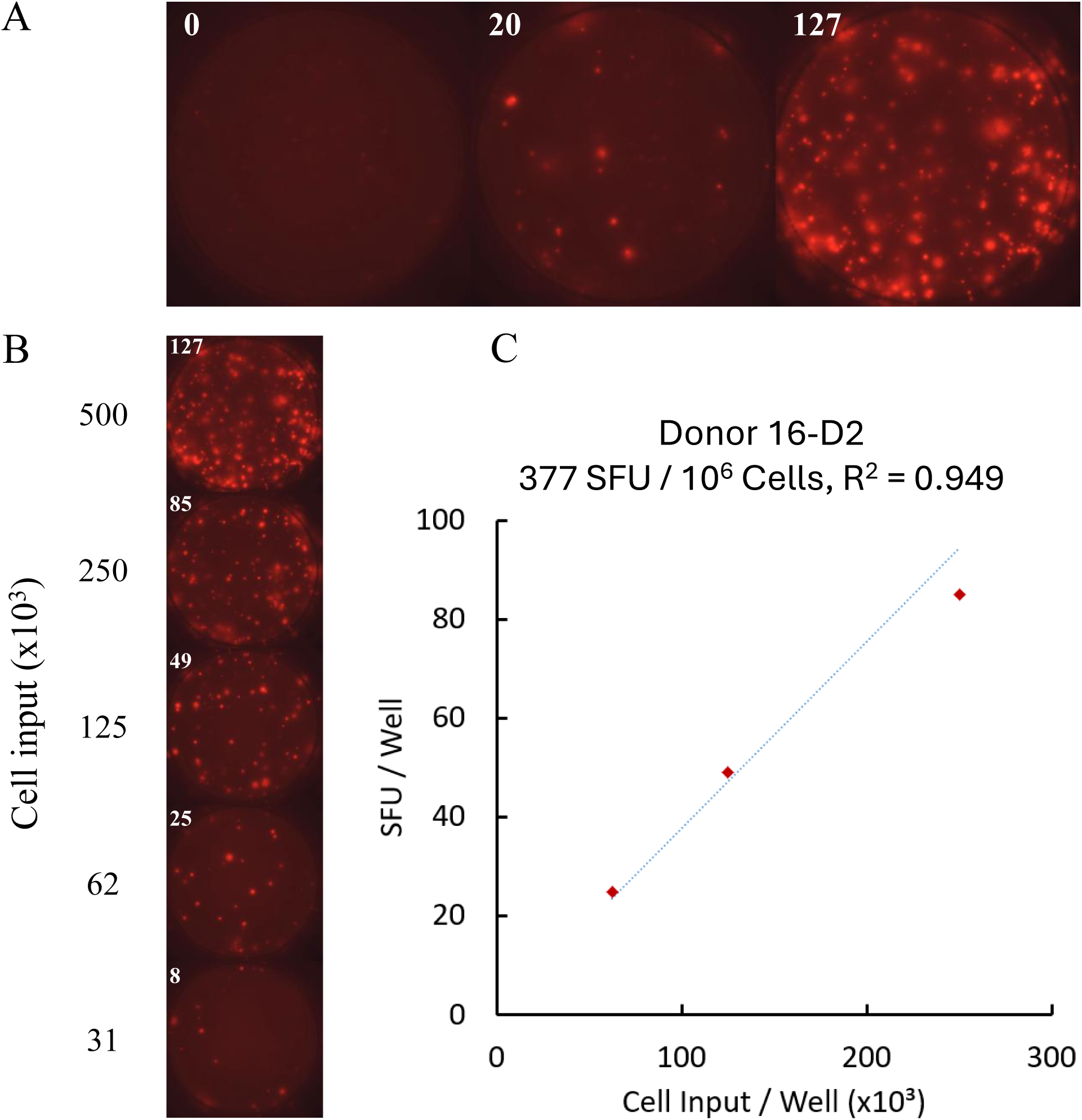
ImmunoSpot® data from a representative subject (Donor-16). (A) Wells from the same donor containing either the pre-vaccination (P) (the well on the left), post-D1 (the middle well) or the post-D2 (well on the right) samples at 5 x 10^5^ PBMC per well. (B) Two-fold serial dilution of the post-D2 bleed sample. The PBMC were seeded in the numbers specified, according to the plate layout shown in Fig. 1B. Secretory footprints generated by S-antigen-specific B_mem_-derived IgG^+^ antibody-secreting cells (ASCs) were visualized as described in *Materials and Methods*. When the secretory footprints (spot-forming units, SFUs) were too crowded at 5 x 10^5^ PBMC/well for accurate enumeration, i.e. was > 70 SFU/well, as was the case for the 2.5 x 10^5^ per well post-D2 sample shown here, frequencies were calculated from the two-fold dilution series as shown in panel (C), and as described in *Materials and Methods*. Data shown in panels (A) and (B) were generated in a single experiment in which the donor’s PBMC were tested in parallel under identical experimental conditions.

**Figure 3.**
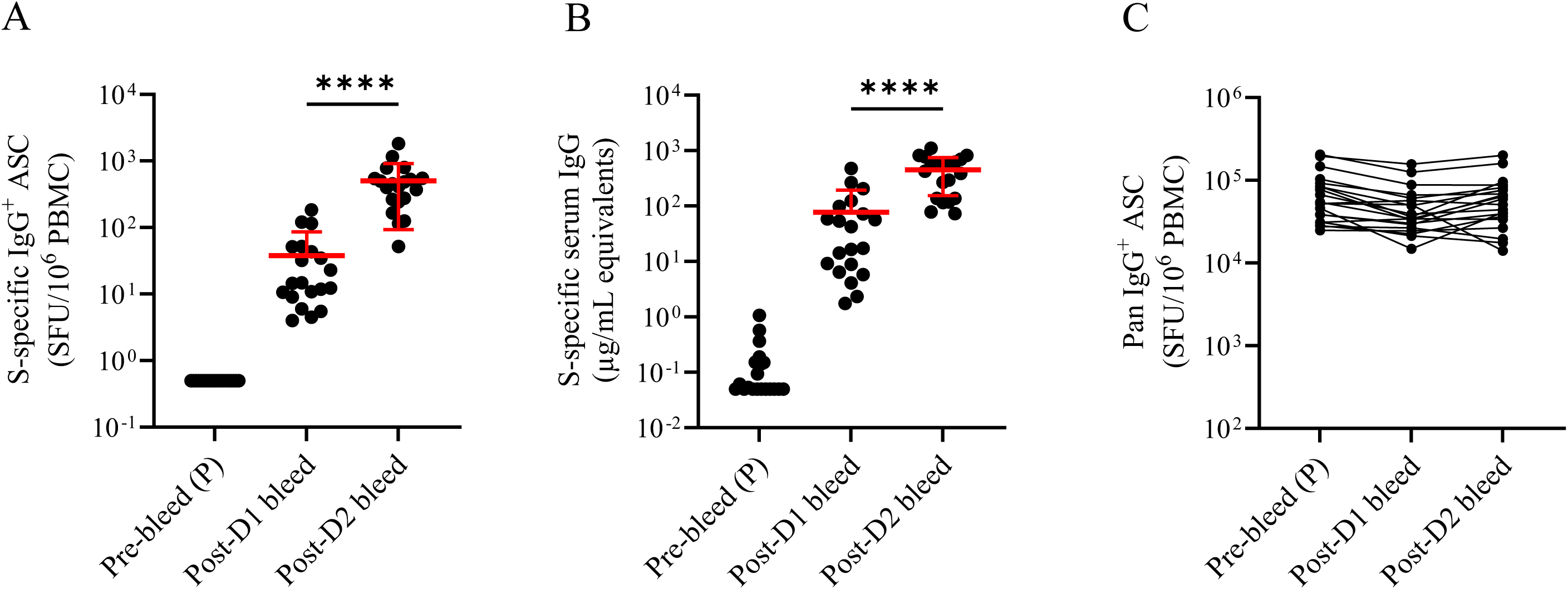
The S-antigen-specific B cell response at the cohort level. (A and B) S-antigen-specific IgG^+^ B_mem_ frequencies (panel A) and IgG serum antibody concentrations with the same specificity (panel B) were measured in the individual donors (each represented by a dot) at the pre-bleed (P), post-D1 and post-D2 timepoints, as specified. To enable visualization on a logarithmic scale the frequency of the S-antigen-specific B_mem_-derived IgG^+^ ASC in the pre-bleed samples was set to 0.5 SFU/10^6^ PBMC, while in reality they were <1 in 2 x 10^6^ PBMC. Likewise, serum IgG antibody levels below our limit of quantification were assigned a value of 0.05µg/mL for graphing purposes. (C) The frequencies of pan (total) IgG^+^ ASC for the three cohorts. The ImmunoSpot® assays for testing the PBMC, and ELISA to test the sera were performed as described in *Materials and Methods*. In panels (A) and (C), the frequencies of S-antigen-specific and pan IgG^+^ ASC were measured in a single experiment in which all PBMC samples were tested in parallel under identical experimental conditions. In panel (B), S-antigen-specific serum IgG levels were measured in two independent experiments and results were combined. The means ± SD are denoted for each cohort in red. Statistical significance between post-D1 and post-D2 timepoints were determined using paired t-tests. ****p<0.0001.

Importantly, reliable B_mem_ measurements by ImmunoSpot® require that PBMC samples be cryopreserved and thawed correctly for the cells to maintain their full functionality (21). As shown in Figure 3C, the overall functionality of the PBMC (*i.e.,* number of pan IgG^+^ ASC detected following polyclonal stimulation) was similar at the pre-bleed, post-D1, and post-D2 timepoints. Therefore, the observed differences in the frequency of S-antigen-specific B_mem_ reactivity was inherent to each donor’s B_mem_ composition at the time of blood collection.

When comparing the relationship between B_mem_ frequencies and IgG titers at the post-D1 timepoint the correlation was weak but already clearly observable (Figure 4A). The dissociation of the humoral B cell response (serum IgG, reflecting plasma cell activity) and its cellular arm, generation of B_mem_, became even more pronounced after the second vaccination (post-D2 timepoint) (Figure 4B). Thus, these data are in line with the notion that serum antibody levels do not reflect on B_mem_ frequencies: i.e. the first and the second walls of B cell-mediated immunity are already dissociated early on during the immune response. But do increased frequencies of B_mem_ predispose to stronger secondary B cell responses, and if not, is the reactivity regulated by antibody feedback?

**Figure 4.**
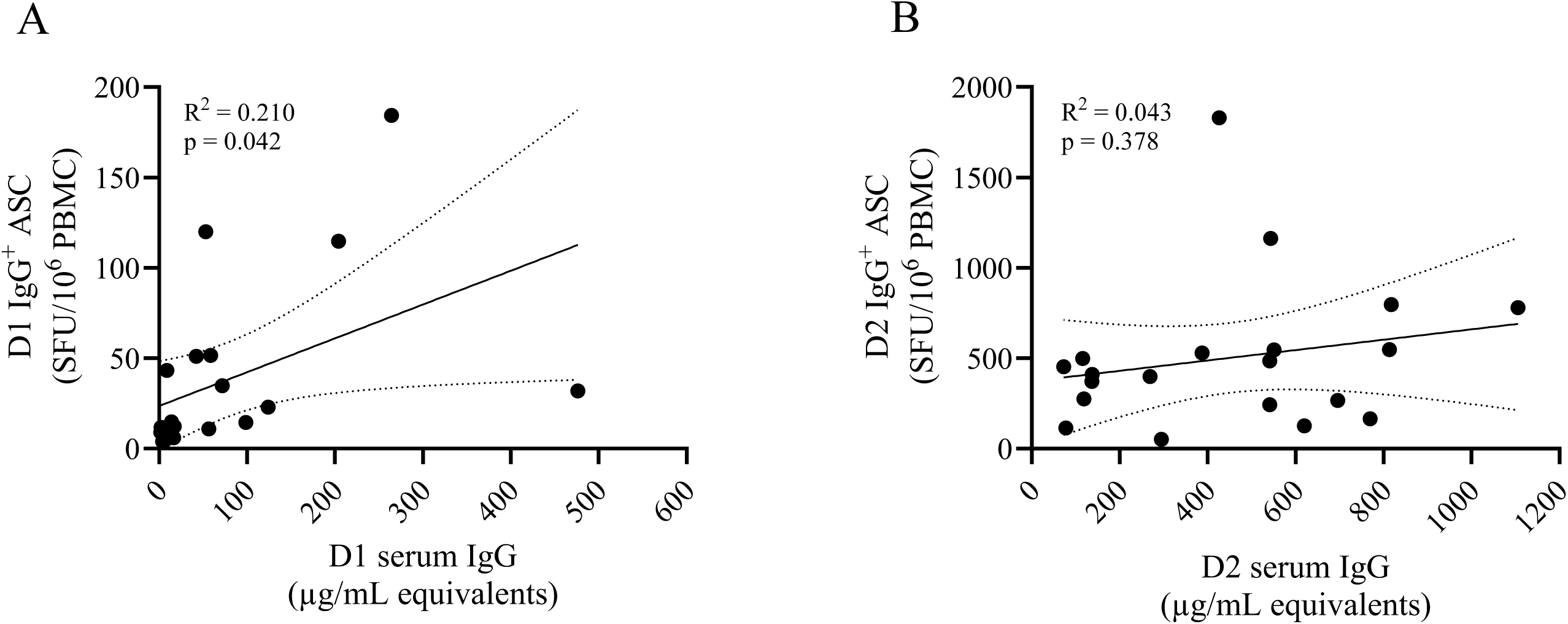
Dissociation of S-antigen-specific serum IgG levels and B_mem_ frequencies in post-D1 (A) and post-D2 (B) bleeds. The results shown in Figure 3 for these two cohorts are broken down here for the individuals with each dot representing one of the vaccinees. Pearson correlation and linear regression analyses were performed. R^2^ and p values are denoted in the corresponding graph.

### 3.3. Relationship between S-antigen-specific serum IgG antibody levels and B_mem_ frequencies

Having measured both arms (humoral and cellular) of the B cell response at the post-D1 and post-D2 timepoints, we next sought to assess how they were correlated. Figure 5A shows that higher numbers of S-antigen-specific IgG^+^ B_mem_ before administering the booster vaccination did not predispose to a higher frequency of B_mem_ at the post-D2 timepoint. Therefore, the presence of a larger antigen-specific B_mem_ pool does not imply a greater degree of recruitment and engagement in a proportionally larger secondary B cell expansion response. Similarly, starting with a greater number of B_mem_ prior to boosting did not lead to a predictably stronger antibody response following the booster vaccination (Figure 5B). Increased antibody levels prior to the boost vaccination also did not predispose to mounting a higher antibody response thereafter (Figure 5C). Collectively, such data suggest that the secondary engagement of antigen-specific precursor cells (i.e. the B_mem_) is not solely driven by the re-exposure to the antigen, but instead is governed by additional regulatory control mechanisms. Indeed, in our donor cohort both the fold increase (clonal expansion) of S-antigen-specific IgG^+^ B_mem_ or the resulting boost in serum IgG levels (*in vivo* differentiation into ASC) was inversely correlated with antibody levels measured at the post-D1 timepoint (Figures 5D and 5E). Consequently, subjects with low levels of S-antigen-specific serum IgG at the post-D1 timepoint exhibited a greater expansion of B_mem_ after the booster vaccination (post-D2).

**Figure 5.**
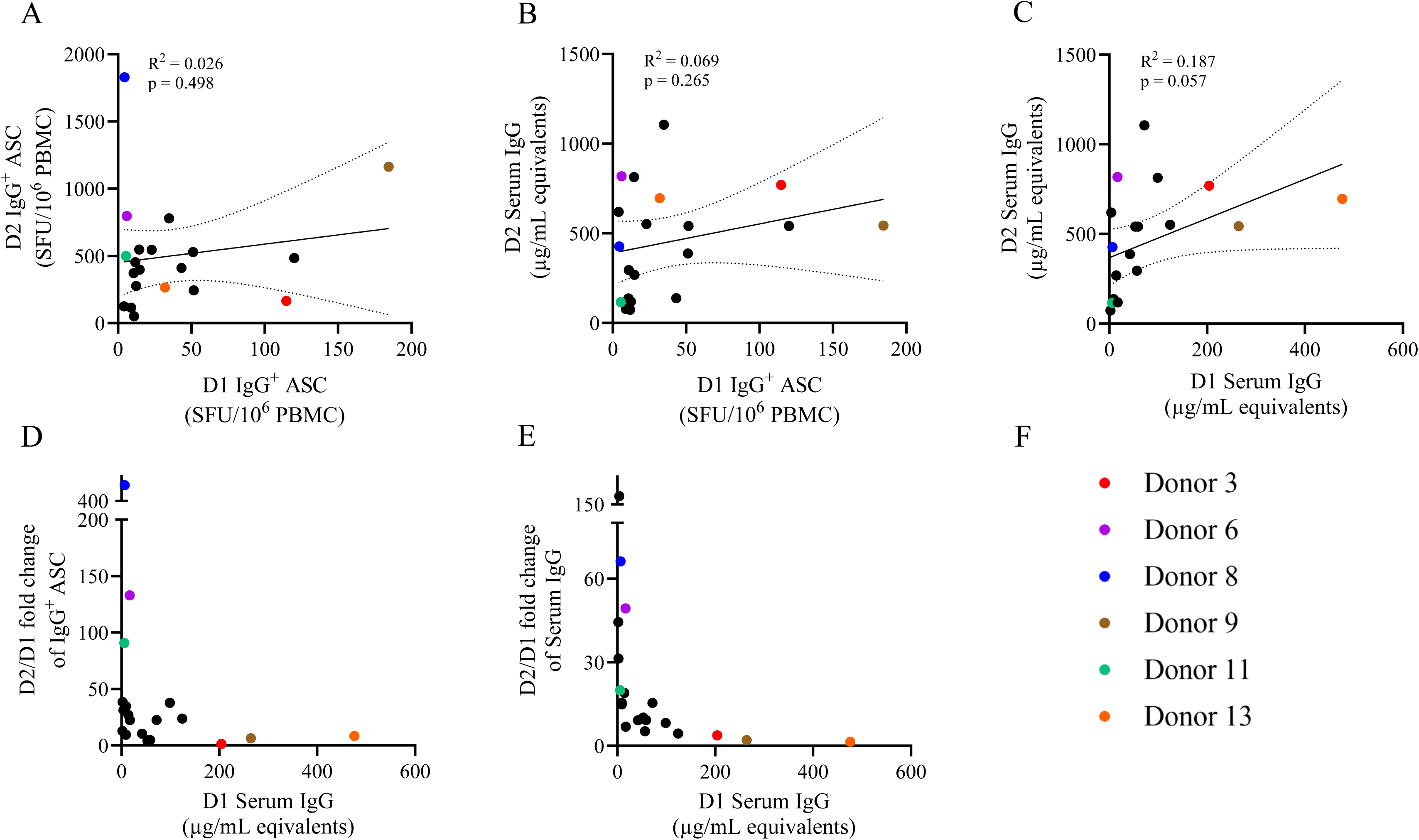
The secondary B cell response to S-antigen negatively correlates with the magnitude of the primary B cell response. The data shown in Figure 3 are represented here for the individual donors, each as a dot, as: (A) post-D1 B_mem_ frequencies on the x-axis vs. post-D2 B_mem_ frequencies on the y-axis, showing that increased B_mem_ precursor frequencies alone did not predispose to proportionally increased B_mem_ frequencies after the booster vaccination. (B) Post-D1 B_mem_ frequencies on the x-axis vs. post-D2 serum antibody levels on the y-axis showing that increased post-D1 B_mem_ frequencies also did not predispose to higher post-D2 serum IgG levels. (C) post-D1 serum IgG levels on the x-axis vs. post-D2 serum IgG levels on the y-axis showing that high antibody levels at the post-D1 timepoint did not predispose to greater antibody levels at the post-D2 timepoint. On the contrary, an inverse relationship is seen: (D) post-D1 serum IgG levels were negatively correlated with the expansion of B_mem_ from post-D1 to post-D2 bleeds, and also with the post-D2 serum antibody increase (E). (F) The identities of six donors of note are depicted by the respective color-coding scheme to enable their tracking in the respective panels. Pearson correlation and linear regression analyses were performed. R^2^ and p values are denoted in the corresponding graphs.

## 4. Concluding Remarks

Adaptive immune responses are based on antigen-driven expansion of specific clones, as originally predicted by Burnet’s clonal selection theory (75). The rationale of vaccinations, including booster administration of the antigen, has been based on this fundamental concept. While correct beyond doubt for primary responses, the assumption has been that the B cell system will stereotypically respond with renewed clonal expansions to each renewed antigen encounter. For the naïve state, when there are by definition no specific antibodies present to exert feedback inhibition, vigorous primary clonal expansions allow antigen-specific naïve B cells (that may be as infrequent as 1 in 10^11^ among all PBMC) to clonally expand to reach frequencies that become detectable, reaching average frequencies of 1 in 10^5^ PBMC (Figure 3A). While the starting number of naïve B cells cannot be known, this might represent as much as a 10-million-fold clonal expansion (corresponding to ∼24 cell division cycles). This initial expansion occurs in the absence of antigen-specific antibodies, and the ensuring primary antibody response becomes detectably soon after (Figure 3B). It is conceivable that the development of the primary S-antigen-specific antibody response within the first week following the initial COVID-19 mRNA (76) can already restrain further clonal expansion of antigen-specific B cells (48). To this end, because the primary serum antibody response in our donor cohort already shows considerable inter-individual variation (Figure 3B), it is plausible that individuals who exhibited a weaker/delayed antibody response engaged in larger B_mem_ clonal expansions and this underlies the divergence in B_mem_ numbers and serum antibody titers observed at the D1 timepoint (Figure 4A).

Overall, while higher antibody levels and B_mem_ frequencies were present after the booster vaccination, the increase is relatively minor relative to the primary B cell response (maximally 400-fold, on average 45-fold, corresponding to maximally 9, on average 5 cell divisions, respectively, Figure 3 and Figure 5). Importantly, this secondary expansion was most pronounced for individuals who failed to develop a robust antibody response after the first vaccine dose (Figure 5D). Conceivably, secondary clonal expansions and the generation of additional waves of B_mem_ and antibody-secreting plasma cells was less inhibited in such individuals following the booster vaccination. Our data suggest that the B cell system does not stereotypically respond to antigen re-encounter with the initiation of a robust secondary B cell response, but does so only when antigen-specific antibodies are not present at sufficiently high levels to be inhibitory. Confirming this notion, vigorous renewed antibody production has been reported in vaccinees in whom serum antibodies have waned (77–79), but for whom unfortunately, B_mem_ data are not available to complete the picture.

In light of emerging data, we contend that it is warranted to re-visit vaccination regimes stipulating the administration of boosters at fixed interval as this assumes that the immune system reacts stereotypically with renewed rounds of clonal expansion and subsequent B_mem_ differentiation following a given antigen stimulus. Indeed, clinical evidence for vaccine attenuation is emerging (80, 81). Extending the immune monitoring data on which clinical vaccination protocols are based, shifting from exclusively titering antibodies to inclusion of cellular measurements (e.g. B_mem_ frequencies and Ig class/subclass usage) is likely to unveil novel insights and guide future vaccination strategies aiming at eliciting durable and protective immunity.

## 5. Limitation of our study

Our study was limited in scope since we only evaluated paired PBMC samples originating from twenty donors. While our cohort included subjects of various ages and both males and females, it lacked racial diversity since only one non-Caucasian participant was included. Additionally, the post-D1 blood draws occurred 2 weeks after the initial vaccination and ∼1 week in advance of the booster vaccination. Therefore, it is plausible that frequencies of S-antigen-specific IgG^+^ B_mem_ and/or serum IgG levels may have continued to increase (or decrease) slightly during this week. Nevertheless, given the differences in response magnitudes observed between individual test subjects in our cohort we consider it highly unlikely that fundamental changes in their response hierarchies would have occurred prior to the administration of their second COVID-19 mRNA vaccine. Likewise, our donor cohort was not profiled for S-antigen-specific serum IgG levels or IgG^+^ B_mem_ frequencies beyond the post-D2 timepoint. Consequently, we were unable to assess the prognostic relationship of serum IgG and B_mem_ frequencies measured acutely after the COVID-19 mRNA vaccine regimen with levels achieved/maintained at longer-term timepoints. Lastly, in contrast to conventional protein subunit/split inactivated vaccines in which the amount of antigen can be measured prior to inoculation, another variable in our donor cohort that could influence the magnitude of the ensuring B cell response was the tissue restriction and duration of S-antigen expression following the COVID-19 mRNA vaccinations (76, 82, 83).

## Supplementary Materials

Supporting information can be downloaded at xxx, Suppl. Figure S1: Illustration of antigen-specific and pan (total) B cell ImmunoSpot® test principles; Suppl. Figure S2: Overview of antibody feedback mechanisms; Table S1: Details of donor cohort.

## Author Contributions

All authors fulfilled the ICMJE recommended criteria for authorship, with their major contribution being as follows: Conceptualization: M.J.H, P.V.L, and G.A.K.; Methodology: N.B., L.Y., and G.A.K; Formal analysis: M.J.H. and G.A.K; Investigation: M.J.H, L.Y., N.B. and G.A.K.; Resources: A.G. and M.T.L.; Data curation: M.J.H., A.G., M.T.L, and G.A.K.; Writing—original draft preparation: PVL; Writing—review and editing M.J.H, S.K., M.T.L., G.A.K., and P.V.L.; Supervision: P.V. L; Project administration: G.A.K. This publication serves as part of M.J.H.’s doctoral thesis to be submitted to University Bonn, Germany. All authors have read and agreed to the published version of the manuscript.

## Funding

This research was funded by the R&D budget of Cellular Technology Limited (CTL).

## Institutional Review Board Statement

PBMC from COVID-19 mRNA vaccinated donors were collected internally at CTL under an Advarra Approved IRB #Pro00043178 (CTL study number: GL20-16 entitled COVID-19 Immune Response Evaluation).

## Informed Consent Statement

Informed consent was obtained from all individuals sampled internally under an Advarra Approved IRB #Pro00043178 (CTL study number: GL20-16 entitled COVID-19 Immune Response Evaluation).

## Data Availability Statement

The data generated in this study will be made available by the authors, without undue reservation, to any qualified researcher.

## Acknowledgments

We thank Graham Pawelec and Alexey Y. Karulin for their valuable discussions and comments on the manuscript. We also want to specifically thank Victoria Gaidaenko from CTL for her help in acquiring PBMC samples.

## Conflicts of Interest

P.V.L. is Founder, President, and CEO of Cellular Technology Limited (CTL), a company that specializes in immune monitoring by ImmunoSpot®. M.T.L. is a cofounder and CSO of CTL. L.Y., N.B., A.G and G.A.K. are employees of CTL. M.J.H. and S.K. declare no conflict of interest. This study was funded by CTL, and the funder directed the study design, collection, analysis, interpretation of data, and the writing of this article, and made the decision to submit it for publication.

## Abbreviations

B_mem_: memory B cell
ASC: antibody-secreting cell
PC: plasma cell
PBMC: peripheral blood mononuclear cell(s)
CM: complete medium
SFU: spot-forming unit
ELISPOT: enzyme linked immunospot assay

