## Supplementary Materials for "The magnitude of the secondary B cell response is primarily defined by antibody feedback inhibition rather than the number of memory B cells present"

**Supplementary Table S1. Details of donor cohort**

| Donor ID | Age | Race | Gender | Vaccine type | Predose sample collection date | Post-D1 bleed sample collection date (14 days after first vaccination) | Post-D2 bleed sample collection date (14 days after second vaccination) |
|---|---|---|---|---|---|---|---|
| Donor 1 | 55-59 | White | M | Pfizer/BioNTech | 4/5/2021 | 4/23/2021 | 5/14/2021 |
| Donor 2 | 35-39 | White | M | Pfizer/BioNTech | 4/2/2021 | 4/19/2021 | 5/10/2021 |
| Donor 3 | 18-19 | White | M | Pfizer/BioNTech | 4/28/2021 | 5/17/2021 | 6/7/2021 |
| Donor 4 | 18-19 | White | M | Pfizer/BioNTech | 3/29/2021 | 4/19/2021 | 5/17/2021 |
| Donor 5 | 60-65 | White | F | Pfizer/BioNTech | 3/15/2021 | 4/5/2021 | 4/26/2021 |
| Donor 6 | 30-34 | White | F | Pfizer/BioNTech | 4/1/2021 | 4/21/2021 | 5/11/2021 |
| Donor 7 | 30-34 | White | F | Pfizer/BioNTech | 3/30/2021 | 4/16/2021 | 5/4/2021 |
| Donor 8 | 50-54 | White | F | Pfizer/BioNTech | 3/15/2021 | 4/5/2021 | 4/26/2021 |
| Donor 9 | 30-34 | Multi | M | Pfizer/BioNTech | 4/8/2021 | 5/3/2021 | 5/24/2021 |
| Donor 10 | 60-65 | White | F | Pfizer/BioNTech | 3/10/2021 | 3/25/2021 | 4/15/2021 |
| Donor 11 | 55-59 | White | F | Pfizer/BioNTech | 3/18/2021 | 4/6/2021 | 4/26/2021 |
| Donor 12 | 25-29 | White | F | Pfizer/BioNTech | 3/30/2021 | 4/16/2021 | 5/5/2021 |
| Donor 13 | 25-29 | White | M | Pfizer/BioNTech | 4/5/2021 | 5/4/2021 | 5/26/2021 |
| Donor 14 | 60-65 | White | F | Pfizer/BioNTech | 3/11/2021 | 3/29/2021 | 4/19/2021 |
| Donor 15 | 25-29 | White | F | Pfizer/BioNTech | 3/29/2021 | 4/12/2021 | 5/3/2021 |
| Donor 16 | 50-54 | White | M | Pfizer/BioNTech | 4/9/2021 | 5/4/2021 | 5/25/2021 |
| Donor 17 | 20-24 | White | F | Pfizer/BioNTech | 4/5/2021 | 4/21/2021 | 5/11/2021 |
| Donor 18 | 50-54 | White | F | Pfizer/BioNTech | 3/17/2021 | 4/23/2021 | 5/14/2021 |
| Donor 19 | 25-29 | White | M | Pfizer/BioNTech | 3/29/2021 | 4/13/2021 | 5/3/2021 |
| Donor 20 | 30-34 | White | M | Pfizer/BioNTech | 4/1/2021 | 4/23/2021 | 5/21/2021 |

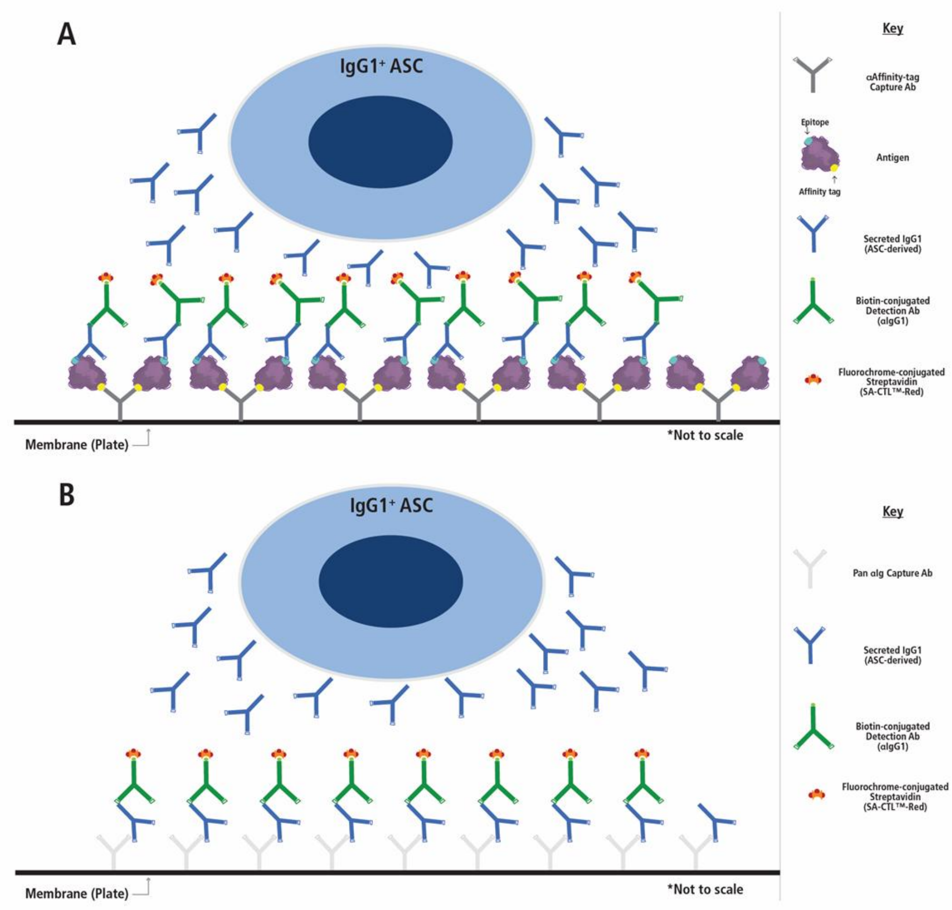


Supplementary Figure S1. Illustration of antigen-specific and pan (total) B cell ImmunoSpot® test principles. A) In an antigen-specific detection assay the membrane can be coated directly (not shown) or via affinity capture (as shown) with the antigen of interest to achieve a high density of antigen coating and maximal detection of antigen-specific antibody-secreting cells (ASCs) [28]. When ASCs are plated onto such an antigen-coated surface, the antibodies produced by an antigen-specific ASC (shown in blue), as opposed to antibodies produced by ASCs specific for other antigens, will be retained on the membrane in close proximity to where the ASC resided on the membrane as a secretory footprint. B) Alternatively, in a pan (total) IgG detection assay the Ig produced by ASCs is captured irrespective of its antigen specificity by an anti-species antibody coated onto the membrane (e.g. goat anti-human Igκ/λ) in close proximity to where the ASC resided on the membrane instead of being secreted into the culture supernatant. In both (A) antigen-specific or (B) pan IgG1 ImmunoSpot® assays, plate-bound IgG1-derived secretory footprints are visualized using an anti-human IgG1-specific detection antibody (depicted in green), followed either by deposition of a precipitating visible substrate in an ELISPOT (not shown) or through selective excitation of the fluorophore associated with the detection antibody and visualization using a suitable instrument in a FluoroSpot (as shown).

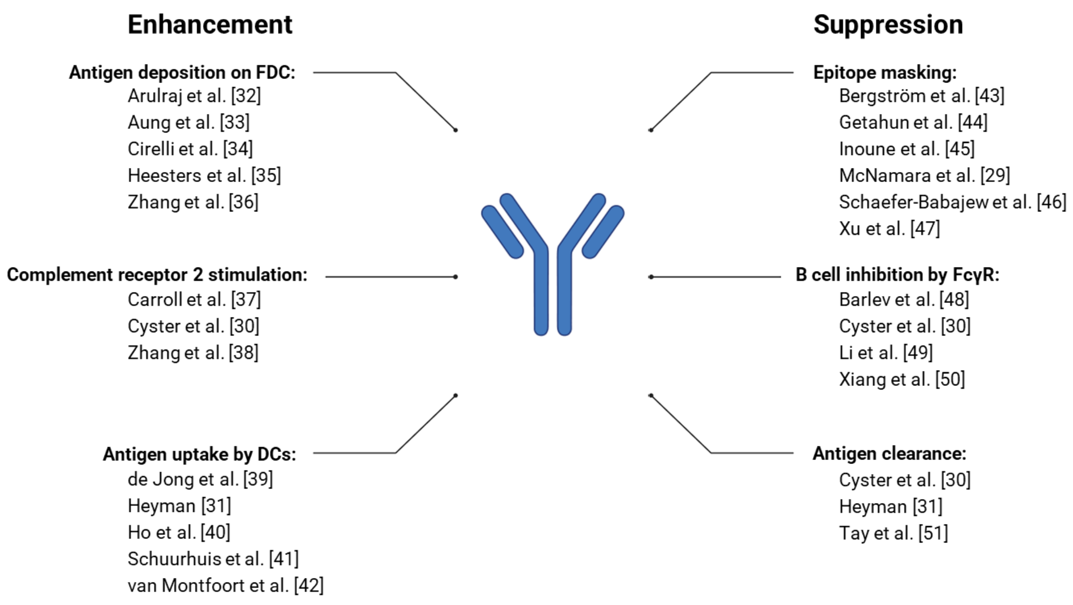


Supplementary Figure S2. Overview of antibody feedback mechanisms. References detailing each mechanism are denoted and numbering of references corresponds to the main text (illustration of antibody originates from BioRender).
